# Patterns of recurrence after curative-intent surgery for pancreas cancer reinforce the importance of locoregional control and adjuvant chemotherapy

**DOI:** 10.1101/270884

**Authors:** Rohan Munir, Kjetil Soreide, Rajan Ravindran, James J. Powell, Ewen M. Harrison, Anya Adair, Stephen J. Wigmore, Rowan W. Parks, O. James Garden, Lorraine Kirkpatrick, Lucy R Wall, Alan Christie, Ian Penman, Norma McAvoy, Vicki Save, Alan Stockman, David Worrall, Hamish Ireland, Graeme Weir, Neil Masson, Chris Hay, James-Gordon Smith, Damian J. Mole

## Abstract

**Introduction:** The pattern of recurrence after surgical excision of pancreas cancer may guide alternative pre-operative strategies to either detect occult disease or need for chemotherapy. This study investigated patterns of recurrence after pancreatic surgery.

**Methods:** Recurrence patterns were described in a series of resected pancreas cancers over a 2-year period and recurrence risk expressed as odds ratio (OR) with 95% confidence interval (C.I.). Survival was displayed by Kaplan-Meier curves.

**Results:** Of 107 pancreas resections, 69 (65%) had pancreatic cancer. R0 resection was achieved in 21 of 69 (30.4%). Analysis was based on 66 patients who survived 30 days after surgery with median follow up 21 months. Recurrence developed in 41 (62.1%) patients with median time to first recurrence of 13.3 months (interquartile range 6.9, 20.8 months). Recurrence site was most frequently locoregional (n=28, 42%), followed by liver (n=23, 35%), lymph nodes (n=21, 32%), and lungs (n=13, 19%). In patients with recurrence, 9 of 41 had single site recurrence; the remaining 32 patients had more than one site of recurrence. Locoregional recurrence was associated with R+ resection (53% vs 25% for R+ vs R0, respectively; OR 3.5, 95% C.I. 1.1-11.2; P=0.034). Venous invasion was associated with overall recurrence risk (OR 3.3, 95% C.I. 1.1-9.4; P=0.025). In multivariable analysis, R-stage and adjuvant chemotherapy predicted longer survival.

**Discussion:** The predominant locoregional recurrence pattern, multiple sites of recurrence and a high R+ resection rate reflect the difficulty in achieving initial local disease control.

## Introduction

Pancreatic adenocarcinoma is one of the most lethal cancers. Survival is reported at less than 10% at 5 years, even after surgery with curative intent and with best available adjuvant chemotherapy^1^. Infrequent early diagnosis, lack of efficacious systemic anti-cancer therapies, and difficulty in achieving a radical resection at the time of surgery contribute to the challenging outlook. Even when clear resection margins are achieved, cancer often relapses early in the postoperative course, with a detrimental outcome. However, it is generally accepted that radical surgical resection with systemic chemotherapy either in a neoadjuvant or adjuvant setting offers the only evidence-based treatment with a chance for cure^2^. Initial staging should classify localised tumours as resectable or unresectable. Notably, the term ‘unresectable’ (or, ‘locally advanced pancreatic cancer’) currently contains a large ill-defined category of potentially resectable disease that is nominated ‘borderline resectable’. Several definitions exist for the term borderline, although an international consensus has been provided^3^. Both surgical and oncological strategies vary between institutions in the protocols for neoadjuvant and adjuvant treatment. With standardised pathology reporting, R+ status of the pathology specimen is the norm, and this fact supports the rationale for neoadjuvant therapy.

Regardless of the variations in definitions of resectability based on multidisciplinary evaluation of imaging studies or post-operative clear margins on pathology, the clinical information on recurrence patterns in resected pancreas cancers may give information on the natural progression of disease after surgery. It may further give information for the rationale for additional preoperative staging that may help patient selection for surgery or neoadjuvant therapy (e.g. MRI to detect isolated occult liver metastasis) or give clues as to where disease control fails, i.e. whether locoregional or systemic.

The aim of this study was therefore to examine the proportion of patients and patterns of pancreas cancer recurrence to contribute baseline data that may assist in the future design of staging and treatment pathways.

## Methods

### Ethics

This study was classed as audit and therefore exempted from formal Research Ethics Committee review. Caldicott Guardian approval was sought and granted.

### Study design

This was a retrospective observational cohort of all major pancreatic resections for suspected malignancy between 1st Jan 2012 and 31st Dec 2013 at the Royal Infirmary of Edinburgh, UK.

This observational study is reported according to the STROBE guidelines^4^, as best applicable.

### Inclusion and exclusion criteria

All pancreatic resections during the period were included. Patients who had benign diseases, secondary malignancies, pancreatic neuroendocrine tumours (pNETs) and rare tumours were excluded from the study question on recurrence pattern. Patients who died within 30 days were excluded from recurrence analysis, as no relevant follow-up time could be used for evaluation. Thus, primary malignancies of the pancreas included for recurrence patterns were pancreatic ductal adenocarcinoma (PDAC) and distal cholangiocarcinoma (CCA).

### Patient management

All patients with a suspected malignant tumour of the pancreas underwent contrast-enhanced dual-phase multidetector computed tomography (CT) and were discussed in a multidisciplinary team (MDT) meeting, involving surgeons, radiologists and oncologists. No patients in this series underwent neoadjuvant chemotherapy and all patients considered for inclusion were regarded as having surgery with curative intent for cancers within the pancreas. Gross specimen and histopathology were reported by trained pathologists according to recommendations set out by Verbeke et al.^5^. Adjuvant chemotherapy decisions were discussed in the MDT and based on histopathology (pancreatic adenocarcinoma versus cholangiocarcinoma), patient overall fitness, Karnofsky performance score and patient preference. Follow-up included clinical visits and imaging (predominantly CT) as deemed indicated, usually according to symptoms or related to adjunct therapy, and was not subject to a formal protocol.

### Data collection

All data were collected by the first author (RM), with survival data updated in December 2017 by the corresponding author (DJM), and chemotherapy data verified by authors AC and LRW in Jan 2018. Demographics details, pre-operative staging, operative details, pathological staging, morbidity and mortality data were linked with surveillance imaging and clinical follow-up data to define patterns of cancer recurrence. All patients underwent pancreatic surgery with curative intent (Whipple pancreaticoduodenectomy, distal pancreatectomy or total pancreatectomy).

### Outcomes

The primary outcome of interest was the anatomical location of cancer recurrence and its associated factors. Additionally, survival of patients was analysed by recurrence, pathological staging and systemic anti-cancer therapy.

## Statistics

Statistical analyses were done using Statistical Package for Social Sciences (SPSS) for Mac (IBM^®^ SPSS^®^ Statistics, v. 24). Continuous variables were analysed by Kruskal-Wallis assuming a non-parametric distribution and categories were investigated with Chi-square or Fisher’s exact test, as appropriate. Risk was expressed as odds ratio (OR) with 95% confidence interval (C.I.). Survival was displayed by Kaplan-Meier curves and analysed by the log rank test. Statistical significance was set at P < 0.05 and all tests were two-tailed.

## Results

A total number of 107 patients were identified as having undergone a major pancreatic resection, of whom 81 patients had a Whipple’s procedure (76.4%), and the remainder had distal pancreatectomy (n=21, 19.8%) or total pancreatectomy (n=5, 4.7%). Overall operative 30-day mortality was 2.8% and at 90-days was 7.5%.

Based on histopathology, the final diagnosis was non-malignant in 25 patients, and these were thus excluded from the current analysis (**Figure 1**). These included 9 periampullary adenomas, 5 intraductal papillary mucinous neoplasia, 2 chronic pancreatitis, 2 benign cysts not otherwise specified, 1 mucinous cystic neoplasm, 1 cholelithiasis, 1 primary sclerosing cholangitis, and 1 benign neuroendocrine tumour.

**Figure 1.**
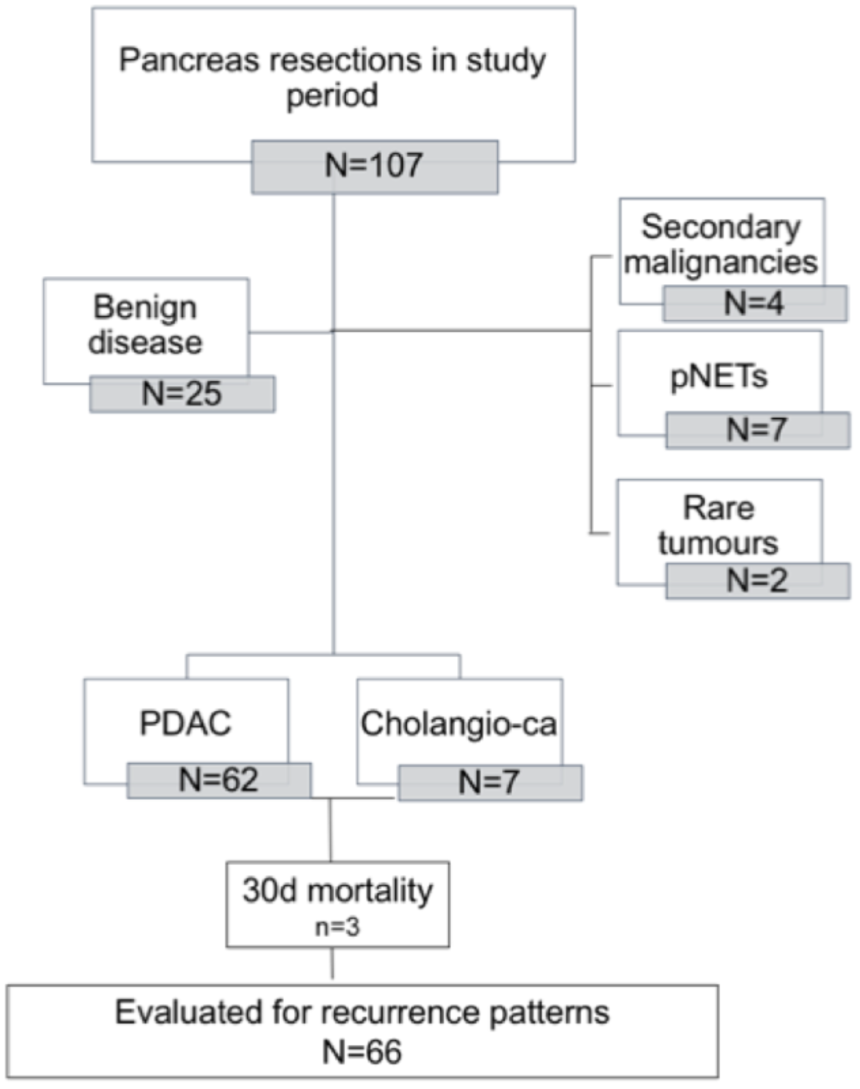
Study flowchart.

Malignant disease was confirmed on pathology in 82 patients with pancreatic ductal adenocarcinoma (PDAC) in 62 patients (76%), distal cholangiocarcinoma (CCA) in 7 patients (8.6%), malignant neuroendocrine tumour in 7 patients (8.6%) and rare tumour types (leiomyosarcoma, renal cell metastases, mixed neuroendocrine and GIST) in the remaining 6 patients (7.3%).

Of 69 patients with PDAC or CCA, there were 3 postoperative deaths which prevented follow up of recurrence (**Figure 1**). For the patients included, the median age was 66 years (range 42-82 years), with 25 (38%) being female. The patients had a median follow-up time after surgery of 20.7 months (interquartile range 37.1 months), and median time from surgery to first radiological imaging was 12.4 months (IQR 6.3 to 22.2 months). There was no difference in follow-up time or time to first scan after surgery in patients with or without overall recurrence (any site), nor specifically for locoregional, liver, lung or lymph node recurrences.

Post-operative chemotherapy was administered in 32 patients (48.5%), given with adjuvant intent in 26 patients (39.4%), palliative intent for early recurrent disease in 5 patients (7.6%), while 1 patient (1.5%) was given post-operative chemotherapy following an R2 resection. Gemcitabine monotherapy was prescribed most frequently. Details of the adjuvant chemotherapy regimens are given in Table 1. Neoadjuvant chemotherapy was not given to any patient in this cohort. No patients received radiotherapy.

**Table 1.** Details of adjuvant chemotherapy

|  | Chemotherapy regimen | Number of patients | % of cohort |
| --- | --- | --- | --- |
| Adjuvant | Gemcitabine | 16 | 24.2 |
|  | Capecitabine | 9* | 13.6* |
|  | Gemcitabine/capecitabine | 2 | 3.0 |
|  |  | 5 | 7.6 |
| No adjuvant chemotherapy |  | 39 | 59.1 |
| Total |  | 66 | 100 |

\* Includes 1 patient who received adjuvant capecitabine following an R2 resection.

### Patterns of recurrence

In 66 patients evaluated, recurrence developed in 41 patients (61%), with a median time to first recurrence of 13.3 months (IQR 6.9, 20.8 months). Two patients died without a documented site of recurrence.

The most frequent site of recurrence was locoregional (n=28), followed by liver (n=23), lymph nodes (n=21), and lung (n=18). Two patients had recurrences in bone and peritoneum (**Figure 2**).

**Figure 2.**
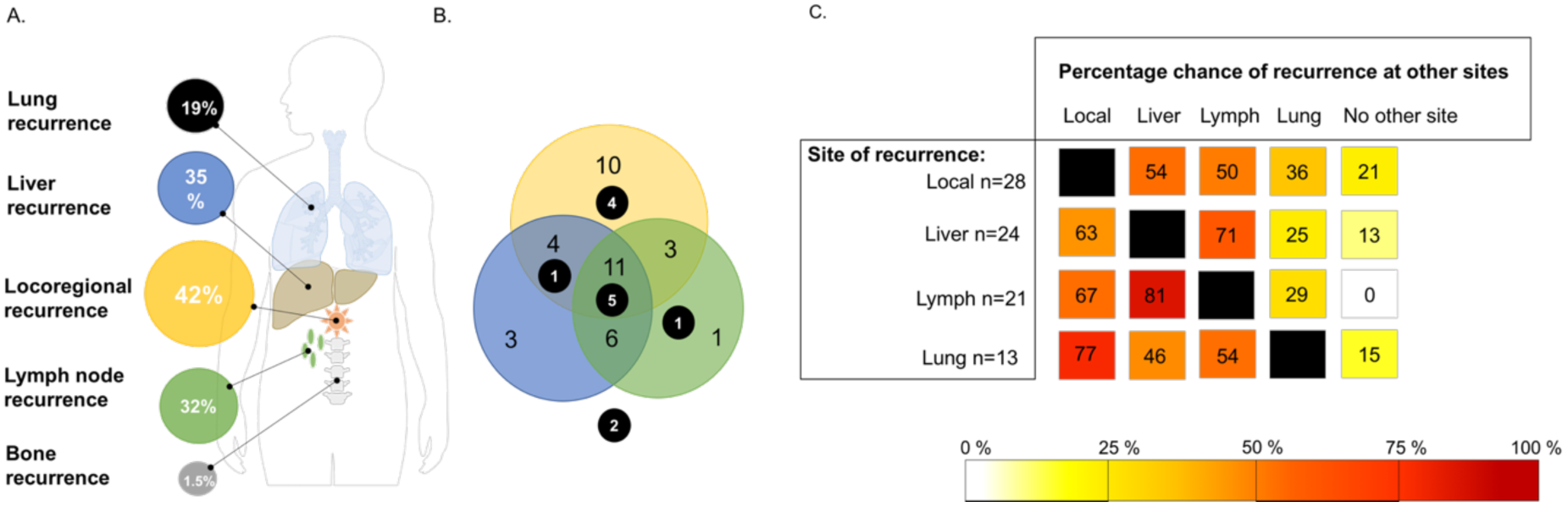
Patterns of disease recurrence. (A) Distribution of metastases according to anatomical site. Each patient may have had more than one sites of recurrence. (B) Modified Venn diagram showing the overlap between single and multiple sites of recurrence. (C) Risk of recurrence at other sites given a defined anatomical site of recurrence.

The majority of liver recurrence (15 of 23) occurred in patients with locoregional recurrences rather than as isolated liver metastasis (OR 3.3, 95% C.I. 1.2-9.6; P=0.024). Locoregional recurrence was also present in the majority (14 of 21) with lymph node recurrence (OR 4.0, 1.3-12.2; P=0.012). Only 3 patients had liver-only as first recurrence site.

R0 resection was achieved in 21/69 (30.4%). A documented R+ resection was associated with higher risk (23 of 43) for locoregional recurrence compared to R0 (5 of 20) resections (53% vs 25%, respectively. OR 3.5, 95% C.I. 1.1, 11.2; P=0.034), but not for overall recurrence, nor recurrence to liver, lung, lymph nodes or other sites.

Presence of histological venous invasion was significantly associated with overall recurrence risk (venous invasion positive: 28 of 38 with recurrence, 74%) compared to no venous invasion (12 of 26 with recurrence, 46%), for an OR 3.3, 95% ci 1.1-9.4; P=0.025).

### Recurrence patterns in potentially favourable groups

The potentially favourable patients with T1 disease (n=4) experienced recurrence in the liver (n=2), lungs (n=2), lymph nodes (n=1), and locoregional (n=2), demonstrating metastatic capacity even with small tumour size. Similarly, of the 12 patients who had pN0 after surgery, 5 recurred in liver, 4 in lymph nodes, 4 locoregional and 3 in lungs. Of patients who achieved an R0 resection (n=21), there were 7 liver recurrences, 5 lymph node recurrences, 4 lung recurrences and 5 local recurrences.

### Survival according to recurrence

Mean overall survival (because the median was not reached for R0 patients) was longer after complete pathological resection (R0, 3.54 years (95% C.I. 2.75, 4.32) vs R1, 1.95 years (95% C.I. 1.53, 2.38; Log Rank test P = 0.005) (**Figure 3**). Data from one patient categorised as R2 were excluded from the survival analysis to allow a simpler two-group survival analysis. The leading factor for survival was being recurrence free or not (**Figure 4A**). Among those with recurrent disease, there was no survival difference according to the anatomical site of recurrence being in liver, lymph nodes or locoregional (**Figure 4B, C, D**). Notably, most patients had recurrence at more than one site.

**Figure 3.**
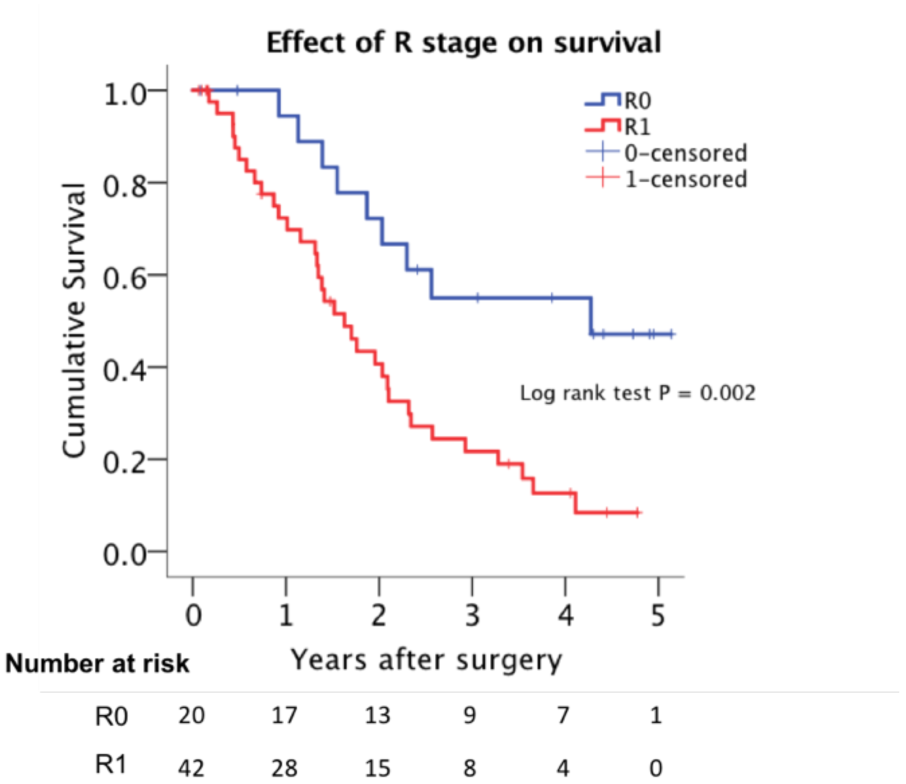
Survival by resection margin status.

**Figure 4.**
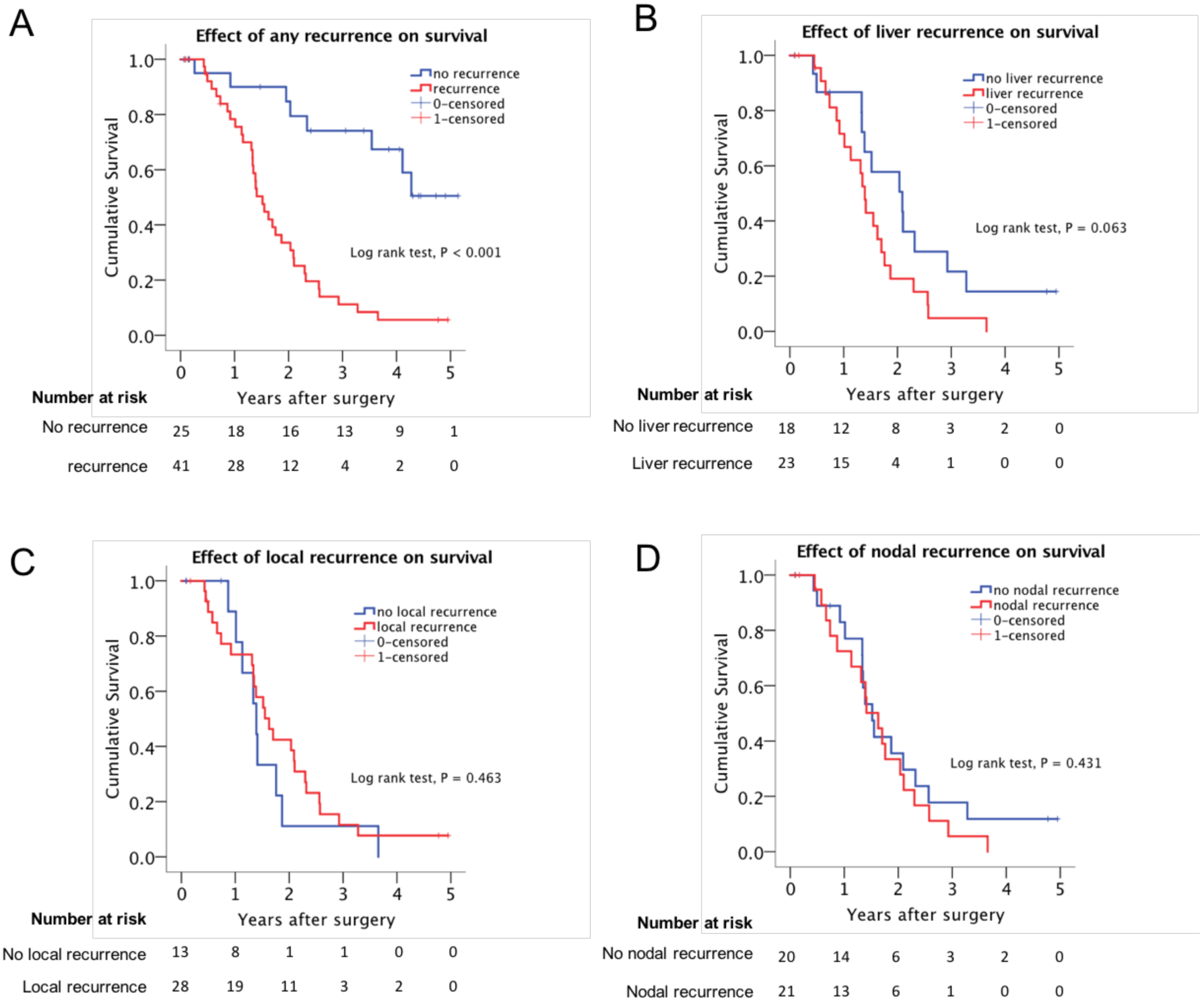
Survival by anatomical site of recurrence.

Overall survival for patients who received adjuvant chemotherapy was longer, but in our cohort this benefit was only seen in those with R1 resection margin status. (**Figure 5A and B**). Median survival was 2.75 years (95% C.I. 2.12, 3.39) in patients with R1 resections, if chemotherapy was given, versus 1.22 years (95% C.I. 0.91, 1.54) if chemotherapy was not given (Log Rank test P < 0.001).

**Figure 5.**
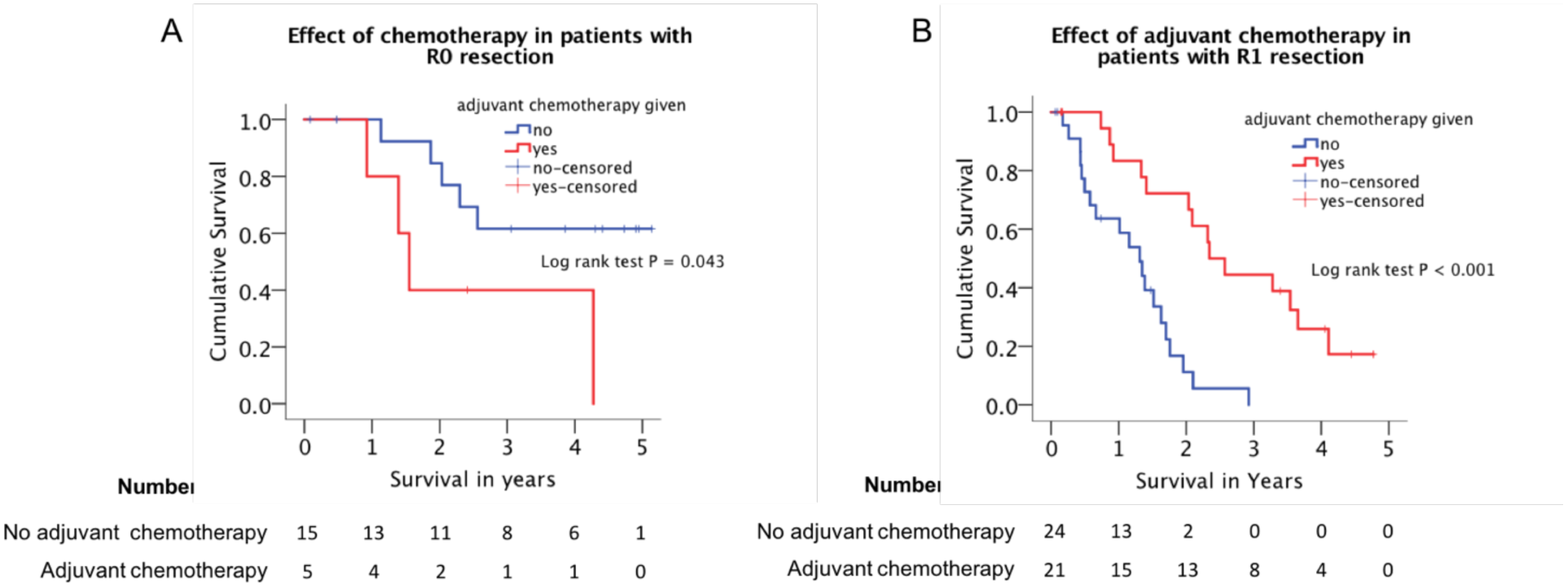
Survival association of chemotherapy.

## Discussion

This observational study demonstrates the recurrence patterns in patients with surgically resected pancreatic cancer in a group of patients to whom neoadjuvant chemotherapy or radiotherapy was not administered. The observed pattern of recurrence was predominantly locoregional, followed by several distant metastatic sites. This was observed despite the fact that locoregional recurrence can be difficult to detect unequivocally on imaging. The time to recurrence was approximately one year, with a considerable number recurring earlier in the post-operative course. Although it is acknowledged that the number of patients in this cohort is small to draw strong conclusions, in the subgroups with potentially favourable features including small tumours (T1), radical resections (R0) and node negative disease (pN0), there were recurrences, both locoregional and distant metastases at multiple sites. Thus, this study corroborates pancreatic cancer as a biologically aggressive disease for which surgery alone is unlikely to cure the disease for the great majority of patients. While local control through negative resection margins is reaffirmed as the most important determinant of survival, and local recurrence is the dominant pattern, the systemic nature of this disease is manifested by the frequent development of metastases in liver, lungs and lymph nodes. Other studies have reported recurrence patterns in PDAC and CCA that have higher rates of distant metastases, rather than locoregional, as site of first recurrence^6, 7^. This may stem from the selected reporting in patients who had a R0 resection (for which locoregional recurrence may be less likely), while the current study reports recurrence for patients with R+ disease. Nevertheless, larger series are needed to establish more definite patterns of recurrence, particularly for these subgroups of patients.

Due to its inherent poor biology, some centres have incorporated more extensive imaging (e.g. MRI and PET-CT) in preoperative staging algorithms to detect disseminated disease prior to resection. However, additional gain from MRI over CT has not been shown to be of particular value in preoperative selection for surgery ^8, 9^. Based on the findings in the current study, additional MRI in the preoperative staging to detect occult liver metastases would have been unlikely to have discovered additional disease that would have prevented progression to surgery. Rather, other markers indicating unfavourable biology may be necessary for directing that phase of decision-making. More importantly, a better patient-centred systemic approach to control of the disease should be the focus for improving survival in patients with pancreatic cancer. Indeed, suggestions of a more aggressive neoadjuvant approach with chemotherapy to most, if not all, patients with resectable disease has been proposed^10-12^. While uncontrolled case-series of neoadjuvant chemotherapy has demonstrated an effect on recurrence patterns in resectable disease^11^, with reduction in both time to as well as number of locoregional and distant metastasis, results from randomised trials are pending for this strategy ^13^.

Surveillance after surgery for pancreatic cancer is controversial. In one study^14^, more patients were detected with asymptomatic recurrence, of which a higher number of patients received further multimodal treatment. Asymptomatic patients thus had longer survival than symptomatic patients^14^. However, there may be a lead-time bias in the difference between the groups, so no clear association can be made on the impact of survival.

The current observational study over a 2-year period is comparable to similar cohorts. A large number of pancreatic resections in this series were performed for other indications than PDAC or CCA. This is corroborated when compared to other series, with sometimes unexpected or benign findings^15^, thus there is a perceived generalisability of the current study population to other settings. However, limitations include a somewhat small cohort which prevents definitive subgroup analysis. Initially, the cohort was explored primarily for the interest of investigating isolated liver recurrences, which were not found as a frequent problem in the series. In the current series, results from the tumour marker CA 19.9 were not available, that may have declared unfavourable disease in some patients^16^. Attempts to further explore small subgroups or more granular details of clinical or pathological factors was not made, to avoid the risk of introducing type II errors and multiple testing errors. This study was not powered for survival analyses, and the survival curves should merely be regarded as descriptive and not explanatory.

The core question in pancreas cancer remains how to best gain locoregional and systemic control of the disease. A better understanding of tumour biology and factors that are amenable to manipulation is desperately needed to improve cancer outcomes. Increased knowledge of the molecular mechanisms in pancreatic cancer is emerging^17^, but has yet to see an impact on clinical management. Better strategies to define what constitutes resectable disease based on pre-operative imaging may include evaluation of initial responses to chemotherapy in selected candidates.

